# Sexual dimorphism of complement-dependent microglial synaptic pruning and other immune pathways in the developing brain

**DOI:** 10.1101/204412

**Authors:** Daria Prilutsky, Alvin T. Kho, Ariel Feiglin, Timothy Hammond, Beth Stevens, Isaac S. Kohane

## Abstract

Sexual dimorphism has been reported in the prevalence, onset and progression of neurodevelopmental and neurodegenerative disorders. We hypothesize that immunological signaling in the developing brain, notably the complement cascade underlying microglial synaptic pruning, could be one mechanism for this dimorphism. Here we show that genes differentially expressed between male and female normal cortical development are enriched for pathways associated with the activation of the innate immune system, complement cascade and phagocytic processes. Specifically, the male brain is enriched for the expression of genes associated with phagocytic function of microglia through complement-dependent synaptic pruning especially at the developmental stages before birth. Our results suggest the existence of a common regulatory module involved in both prenatal immune activation in males and postnatal immune activation in females. The activation of immune pruning pathways at different stages of normal male and female development could provide valuable insights about critical periods of plasticity and refinement in the human cortex that could explain the different vulnerabilities of males and females to neurological disorders.

## Introduction

It has been suggested that there exist sex-based neurobiological differences that either directly promote or increase susceptibility to specific neurological disorders due to certain environmental exposures during development [1]. Sexual dimorphism is present in the prevalence, onset and progression of many neurological disorders, e.g., autism (4:1 male/female), schizophrenia (higher incidence in men), depression (higher incidence in women), Parkinson’s disease (more common in men), and multiple sclerosis (more common in women) [2, 3]. Autism has a particularly striking sex-biased incidence. Recent studies have begun to reveal the molecular mechanisms driving these sexual dimorphisms [4, 5]. Notwithstanding sexual dimorphism, many have reported immunological abnormalities and immune activation in neuropsychiatric disorders such as schizophrenia and autism [6–12]. It is becoming increasingly clear that activation of the brain’s resident immune phagocytic cells, microglia, may contribute to this immune dysregulation [9, 13–16]. Dysfunctional microglia can profoundly affect synaptic development, plasticity and function during neural development. Classical complement cascade, an innate immunity pathway that eliminates pathogens and cellular debris from the periphery, has been implicated in the process of microglial pruning and in tagging synapses for engulfment [17–22]. Complement proteins C1q, C3 and CR3 flag synapses for phagocytosis by microglia and, therefore, function as an “eat-me” signal. Alterations in microglia numbers, morphology and their association with neurons have been noted in autism [14, 15, 23–25]. However, these dysregulated microglial phenotypes have not been directly linked to abnormalities in synaptic pruning. On the other hand, given the importance of maintaining the appropriate neuronal connectivity and evidence of microglia effect, alterations in microglial processes might significantly affect behavior and cognition. Therefore, abnormalities in synaptic pruning may be involved in behavioral phenotypes through attenuated or excess activity (over or under expression) of the complement cascade. Sexual dimorphism in microglial function could therefore partly explain differences in susceptibilities and outcomes of neurological disorders particularly ones arising at key sensitive points of brain development.

Sexual dimorphism in the pattern of microglial colonization and morphology of the developing rodent brain has been described [26]: males have more microglia early in development (P4), while females have more microglia with an activated/amoeboid morphology later in development (P30-P60). The amygdala, hippocampus and cortex have been shown to have more microglia with activated phenotype in females than males in P30 rats [1, 26]. In contrast to the males, “active” microglial cells in P60 females were associated with an increased inflammatory gene expression (pro-inflammatory cytokines, markers of microglial activation) within the hippocampus [1]. Most recently, a microglial development gene expression program was observed to be delayed in male relative to female mice, and exposure of adult male mice to lipopolysaccharide (LPS), a potent immune activator, accelerated microglial development [27].

In this study, we focus on characterizing the differences in innate immunity gene expression in brain development of females and males and the possible consequences of this sexual dimorphism in normal brain wiring and synapse pruning. The activation of immune pruning pathways at different stages of normal male and female development could provide valuable insights about critical periods of plasticity and refinement in the human cortex that could explain the different vulnerabilities of males and females to neurological disorders.

We hypothesize that periods of neurodevelopment previously associated with increased localization of microglia (early development in males; later development in females) might also be characterized by dysregulated microglia and complement-dependent functions including neuronal or synaptic remodeling. Elevated numbers of “active” microglia might result in modified microglia-synapse interactions (aberrant frequency of contacts) and aberrant pruning during similar key periods of brain development. We systematically investigated the expression patterns of genes associated with complement-dependent microglial synaptic processes in each sex across different stages of normal neurodevelopment before and after birth.

## Methods

### Human brain transcriptome data processing

A spatio-temporal transcriptome of the developing human brain (Human Brain Transcriptome [HBT]) has been described previously [28] and these data are publicly available in NIH’s Gene Expression Omnibus (http://www.ncbi.nlm.nih.gov/geo) as GSE25219. These samples were profiled on Affymetrix Human Exon 1.0 ST Array and we used the RMA-normalized transcript (gene)-to-sample series matrix for our analysis with log2 scaled signal for transcript expression. We mapped the 17,565 microarray probes to 16,737 unique human Entrez Gene IDs. We focused on neocortex (NCX) samples at 11 developmental stages, cf. Table 1. The NCX encompasses 11 brain compartments collectively referred as the NCX region. For each microarray probe, we computed the sum of coefficients of variance (coefvar) in stages 2-15. If a gene, represented by its Entrez Gene ID, is mapped to more than one microarray probe, then we pick the probe with the minimal sum of coefvar to represent that gene.

**Table 1:** Detailed description of samples used in this study. M, postnatal months; pcw, post-conceptional weeks (prenatal); Y, postnatal years.

| Stage | Description | Age | Number of Male Samples | Number of Female Samples |
| --- | --- | --- | --- | --- |
| 4 | Early mid-fetal | 13pcw-16pcw | 43 | 22 |
| 5 | Early mid-fetal | 16pcw-19pcw | 46 | 18 |
| 6 | Late mid-fetal | 19pcw-24pcw | 31 | 98 |
| 7 | Late fetal | 24pcw-38pcw | 22 | 24 |
| 9 | Late infancy | 6M-12M | 9 | 33 |
| 10 | Early childhood | 1Y-6Y | 21 | 30 |
| 11 | Middle and late childhood | 6Y-12Y | 20 | 11 |
| 12 | Adolescence | 12Y-20Y | 33 | 32 |
| 13 | Young adulthood | 20Y-40Y | 94 | 60 |
| 14 | Middle adulthood | 40Y-60Y | 42 | 22 |
| 15 | Late adulthood | 60Y≤ | 22 | 42 |

### Differential gene expression analysis and pathway-level analysis

Differential gene expression analysis was performed using a linear regression model (lmFit) as implemented in the limma package in R/BioConductor (http://www.bioconductor.org) with a significance criterion of p<0.05 after false discovery rate (FDR) correction. We used DAVID (The Database for Annotation, Visualization and Integrated Discovery, http://david.abcc.ncifcrf.gov) to identify enriched pathways in significant differentially expressed genes (Entrez Gene IDs) at a Fisher exact p-value threshold (EASE score) less than 0.1. We focused on the Kyoto Encyclopoedia of Genes and Genomes (KEGG) pathways for the top 5% (837 genes ranked by adjusted p-value) significant differentially expressed genes between females and males across all neurodevelopmental stages in order to be consistent across different developmental stages.

## Results

### Stage specific neurodevelopmental genes differentially expressed between males and females are enriched for immunologic pathways

To find transcriptomic differences between male and female brain development, we determined genes that were significantly differentially expressed (DEG) in the neocortex compartment between males and females at each prenatal and postnatal neurodevelopmental stage, separately, in the Human Brain Transcriptome Data [28], Supplementary File 1. For each gene, we also computed the difference in average log2 signal between males and females, logFC = average(log2 signal in males) - average(log2 signal in females) (positive value=over-represented in male; negative value=over-represented in female). Next, we identified Kyoto Encyclopedia of Genes and Genomes (KEGG) pathways that were enriched for the male-female DEG set at each neurodevelopmental stage, Figure 1 and Supplemental File 2. We excluded developmental stages 1-3 and 8 from this analysis as they did not have both male and female samples. Overall, the enriched KEGG pathways were related to immunological response, inflammatory processes, infectious diseases (viral, bacterial and parasitic infections), immune diseases, activation of complement cascade, neurodegenerative diseases and cell-cell interactions. Some stage specific highlights:

**Figure 1:**
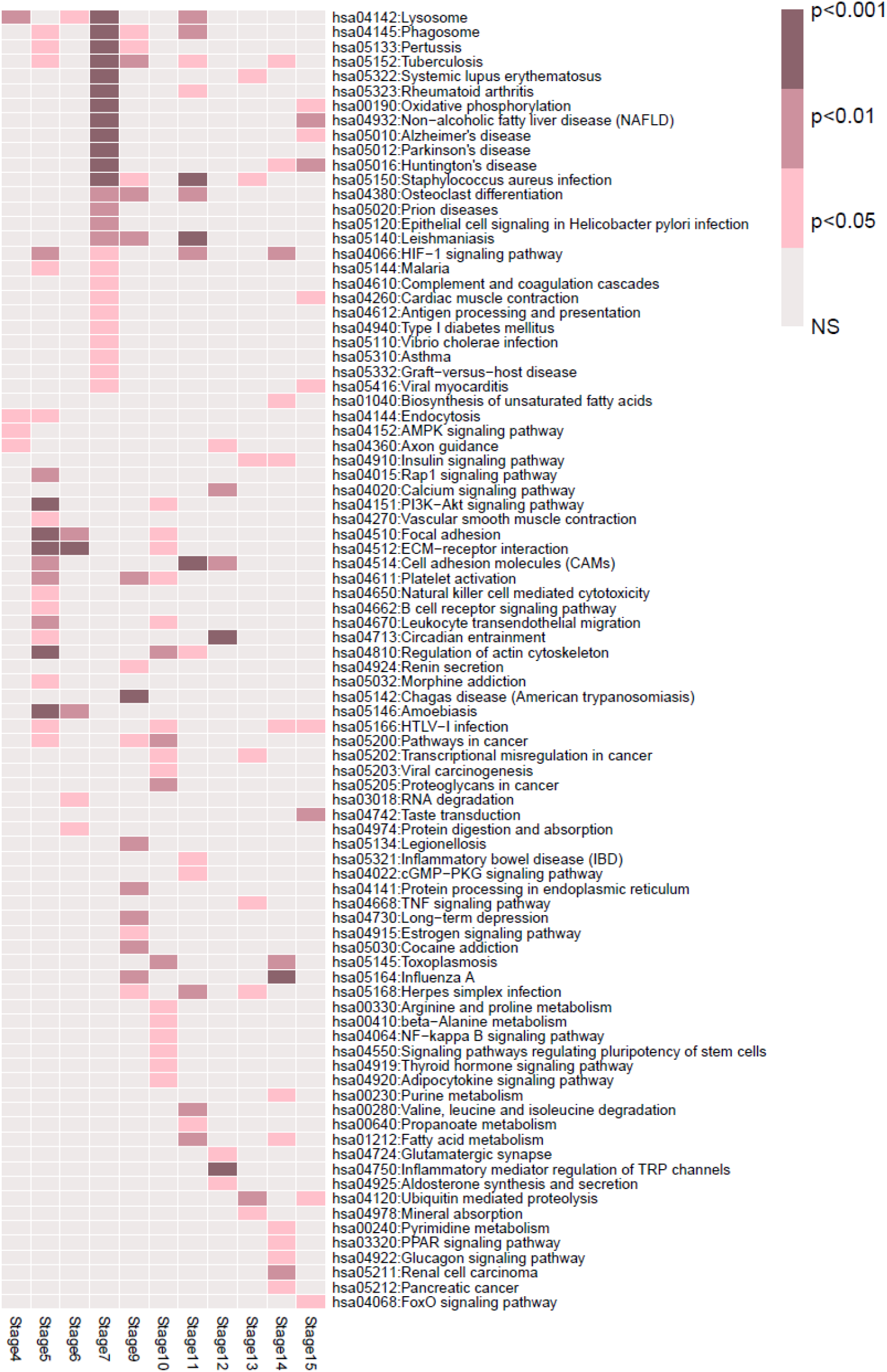
Enriched KEGG pathways in differentially expressed genes in males versus females in cortical brain development per stage. Stages 4–7: Prenatal; Stages 9–15: Postnatal; NS=non-significant.

- <u>*Stage 4*</u>, which corresponds to early-mid fetal brain development is enriched for “Lysosome” (p-value=0.006); “AMPK signaling pathway” (p-value=0.02); “Endocytosis” (p-value=0.03); “Axon guidance” (p-value=0.049).
- <u>*Stage 5*</u>, which corresponds to prenatal early-mid fetal brain development is enriched for cell-cell interactions (“ECM-receptor interaction”, p-value=1.70E-05; “Focal adhesion”, p-value=2.12E-05; “Cell adhesion molecules”, p-value=0.009); parasitic infectious diseases (“Amoebiasis”, p-value=4.96E-05; “Malaria”, p-value=0.03); bacterial infectious diseases (“Tuberculosis”, p-value=0.013; “Pertussis”, p-value=0.03); an interlinked sub-cellular engulfment mechanisms, associated with the process of endocytosis (p-value=0.048) and phagocytosis (“Phagosome”, p-value=0.036); a battery of immune processes (“Leukocyte transendothelial migration”, p-value=0.0018; “Platelet activation”, p-value=0.004; “Natural killer cell mediated cytotoxicity”, p-value=0.016; “B cell receptor signaling pathway”, p-value=0.019).
- At <u>*Stage 6*</u>, which corresponds to late mid-fetal cortical development is enriched for cellcell interactions (“ECM-receptor interaction”, p-value=4.14E-04; “Focal adhesion”, p-value=0.0085).
- <u>*Stage 7*</u>, which corresponds to prenatal late-fetal cortical development that we refer as a “stage before birth” is enriched for bacterial infections (“Staphylococcus aureus infection”, p-value=1.03E-08; “Tuberculosis”, p-value=6.62E-06; “Pertussis”, p-value=1.43E-04; “Epithelial cell signaling in Helicobacter pylori infection”, p-value=0.008773); immune diseases (“Rheumatoid arthritis”, p-value=1.25E-07; “Systemic lupus erythematosus”, p-value=2.51E-04; “Asthma”, p-value=0.019969); neurodegenerative conditions (“Alzheimer’s disease”, p-value=2.04E-07, “Huntington’s disease”, p-value=2.82E-07; “Parkinson’s disease”, p-value=5.84E-05; “Prion diseases”, p-value=0.0015); and activation of immune system, specifically complement system (“Complement and coagulation cascades”, p-value=0.010592; “Antigen processing and presentation”, p-value=0.01923) and phagocytic processes (“Phagosome”, p-value=5.41E-06; “Lysosome”, p-value=6.61E-05). We also performed functional annotation clustering for Stage 7 (Supplementary File 3) and the groups of genes that were common across various over-represented pathways (gene-term association positively reported across pathways) were related to major histocompatibility complexes, interleukin 1 beta, transforming growth factor beta 1, complement system, Fc fragment of IgG receptor.
- <u>*Stage 9*</u>, which corresponds to late infancy (6-12 months of age) that we refer as first “postnatal” stage is enriched for bacterial infections (“Legionellosis”, p-value=0.004066; “Tuberculosis”, p-value=0.005108; “Staphylococcus aureus infection”, p-value=0.014488; “Pertussis”, p-value=0.027557), viral (“Influenza A”, p-value=0.009763; “Herpes simplex infection”, p-value=0.015329) and parasitic infections (“Chagas disease”, p-value=4.40E-04; “Leishmaniasis”, p-value=0.006814).
- <u>*Stage 10*</u>, which corresponds to early childhood is enriched for four major pathway groups: pathways in cancer (“Proteoglycans in cancer”, p-value=0.005145; “Pathways in cancer”, p-value=0.006451; “Transcriptional misregulation in cancer”, p-value=0.035827; “Viral carcinogenesis”, p-value=0.048527), focal adhesion (“Regulation of actin cytoskeleton”, p-value=0.004433; “Focal adhesion”, p-value=0.014427; “ECM-receptor interaction”, p-value=0.042883), immunity (“Platelet activation”, p-value=0.021859; “Leukocyte transendothelial migration”, p-value=0.024017) and endocrine system (“Adipocytokine signaling pathway”, p-value=0.032958; “Thyroid hormone signaling pathway”, p-value= 0.041727).
- <u>*Stage 11*</u>, which corresponds to middle and late childhood (6-12 years of age) is enriched for infections (“Staphylococcus aureus infection”, p-value=9.87E-06; “Leishmaniasis”, p-value=6.90E-04; “Herpes simplex infection”, p-value=0.001805; “Tuberculosis”, p-value=0.014788), immune diseases (“Rheumatoid arthritis”, p-value=0.011647; “Inflammatory bowel disease”, p-value=0.038761) and phagocytic processes (“Lysosome”, p-value=0.002621; “Phagosome”, p-value=0.003777).
- <u>*Stage 12*</u>, which corresponds to adolescence (12-20 years of age) is enriched for neuronal processes (“Calcium signaling pathway”, p-value=0.001017; “Axon guidance”, p-value=0.043581; “Glutamatergic synapse”, p-value=0.049198).
- <u>*Stage 13*</u>, which corresponds to young adulthood is enriched for immune diseases (“Systemic lupus erythematosus”, p-value=0.012957); and viral (“Herpes simplex infection”, p-value=0.026949) and bacterial infections (“Staphylococcus aureus infection”, p-value=0.028174).
- <u>*Stage 14*</u>, which corresponds to middle adulthood (40-60 years of age) is enriched for viral infections (“Influenza A”, p-value=4.89E-04; “HTLV-I infection”, p-value=0.019856; parasitic (“Toxoplasmosis”, p-value=0.002492) and bacterial infections (“Tuberculosis”, p-value=0.017592). Neurodegeneration, specifically “Huntington’s disease” (p-value=0.034474), was over-represented as well.
- <u>*Stage 15*</u>, which corresponds to late adulthood (>60 years of age), was enriched for neurodegenerative conditions (“Huntington’s disease”, p-value=0.004999; “Alzheimer’s disease”, p-value=0.035842) and viral diseases (“HTLV-I infection”, p-value=0.019105; “Viral myocarditis”, p-value=0.039704).

Enriched KEGG pathways within differentially expressed genes across each stage of cortical development are shown in Figure 1. In summary, enriched biological pathways differentiating between male and female cortical development converge on the activation of the immune system.

### Neurodevelopmental genes differentially expressed between males and females before birth converge on the immunological-complement-phagosome axis

We investigated the sex-biased immune-complement-phagosome pathways that were enriched in various neurodevelopmental stages, notably Stage 7, as previous studies have shown these pathways to be associated with dysregulated microglia-mediated synaptic pruning. We omitted Stage 8, referred in the original Human Brain Transcriptome data as neonatal and early infancy stage immediately after birth (0-6 months of age), because it had no female samples. The stage after birth in our analysis is Stage 9. The heatmap in Figure 2 shows the male versus female LogFC (see above result) for gene constituents of the KEGG “Complement and coagulation system” at different neocortical developmental stages. We noted that these genes were broadly more highly expressed in males than females at Stage 7 (before birth), and more highly expressed in females than males at Stage 9 (after birth). Interestingly, we observed that some of the Stage 7 genes were again highly expressed in males than females at Stage 11.

**Figure 2:**
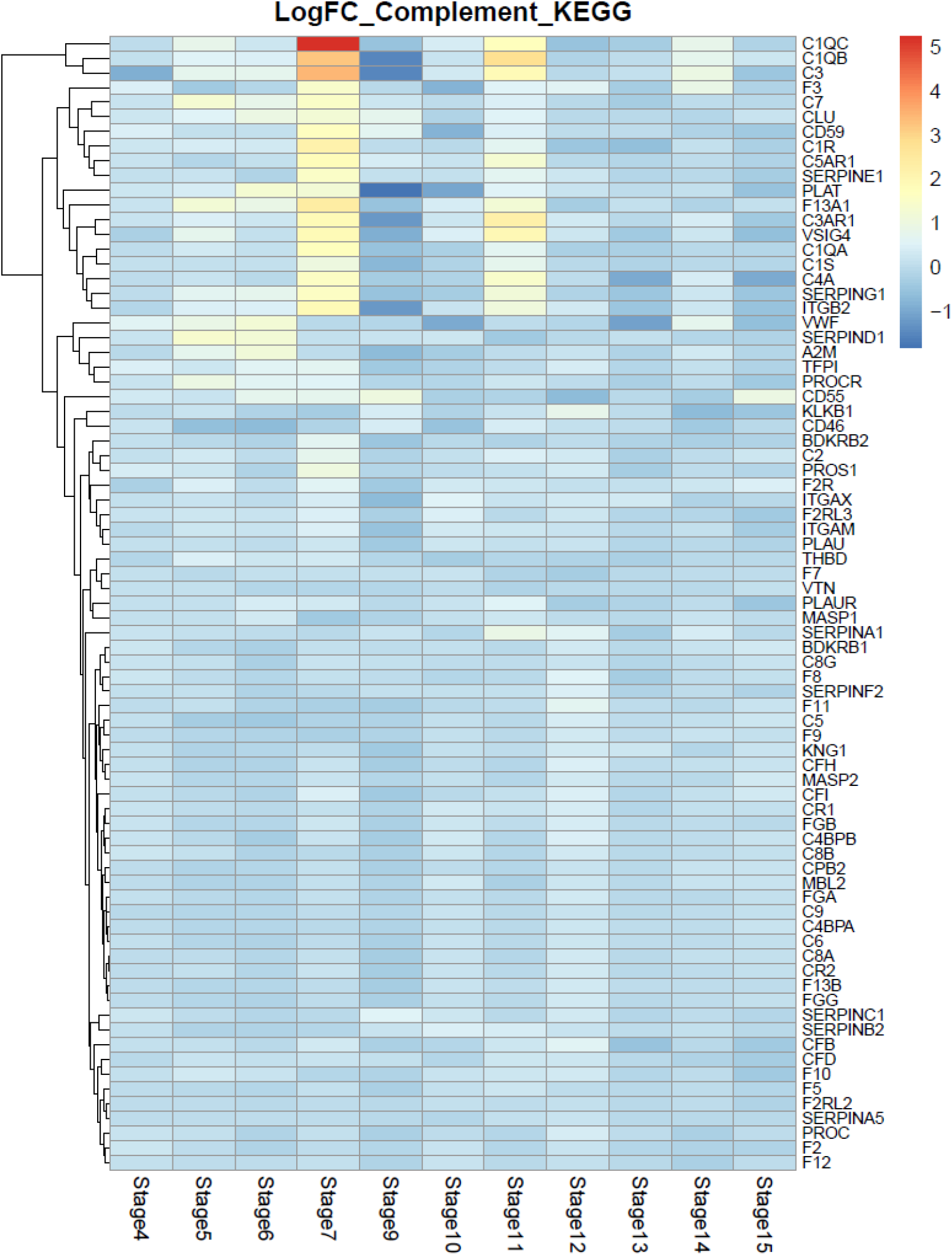
Hierarchical clustering of logFC (Fold Change=Males/Females) of expression of Complement cascade (KEGG pathways) in males versus females. Clustering was performed only on rows (genes). Red=Upregulated in males versus females; Blue=Upregulated in females versus males; Stage 7=Before birth; Stage 9=After birth.

Table 2 shows genes that were significantly (FDR <0.05; logFC >1 i.e., 2-fold) upregulated in the brain tissue in males compared to the females. Notably C1QC, which forms a complex with C1QA (“tags” synapses for elimination)/C1QB, and likely participates in synaptic pruning, is one of the most significantly differentially expressed genes between males and females, ranked #5 out of 16737 based on the adjusted p-value (logFC=5.26, p-value= 2.075E-35). Interestingly, several genes within top 50 differentially expressed genes out of 16737 (Table 2) were related to immunological signaling in developing brain notably:

**Table 2:**
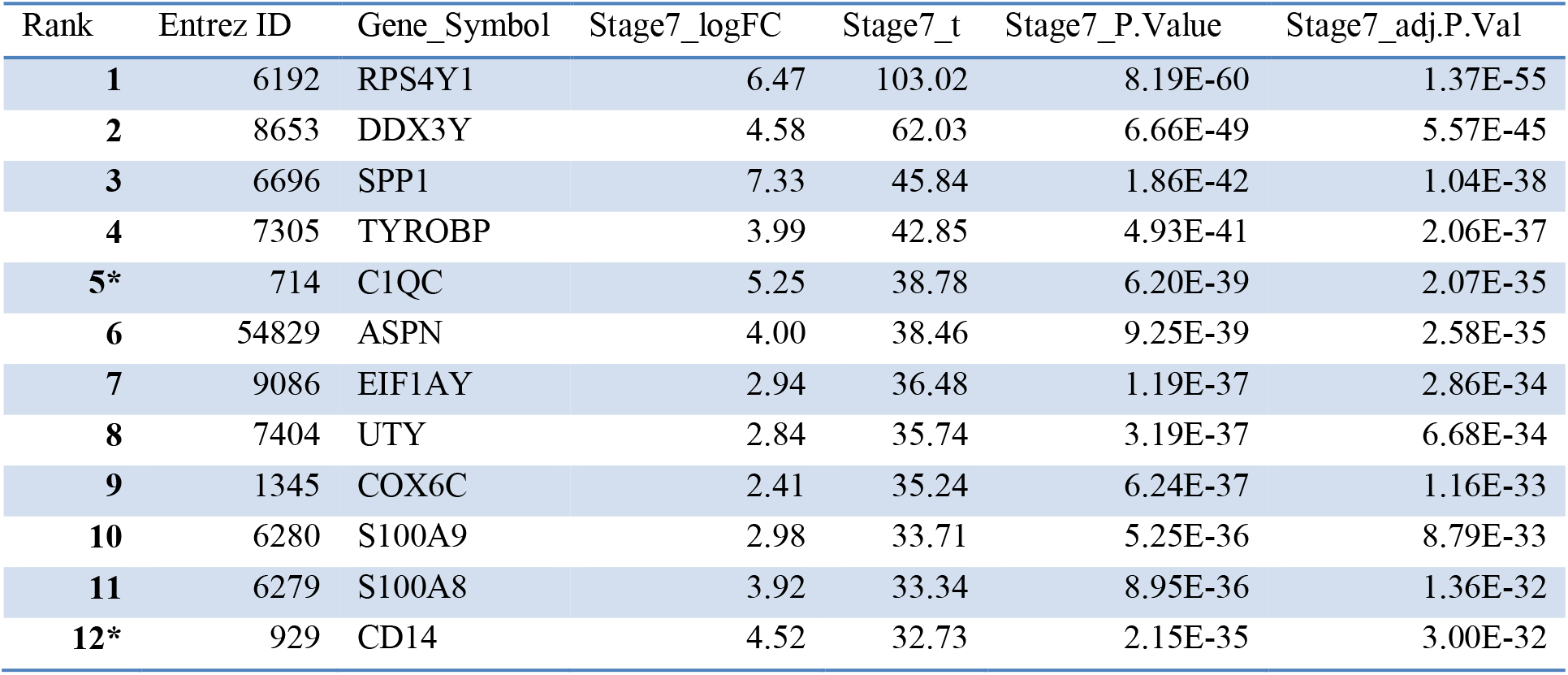
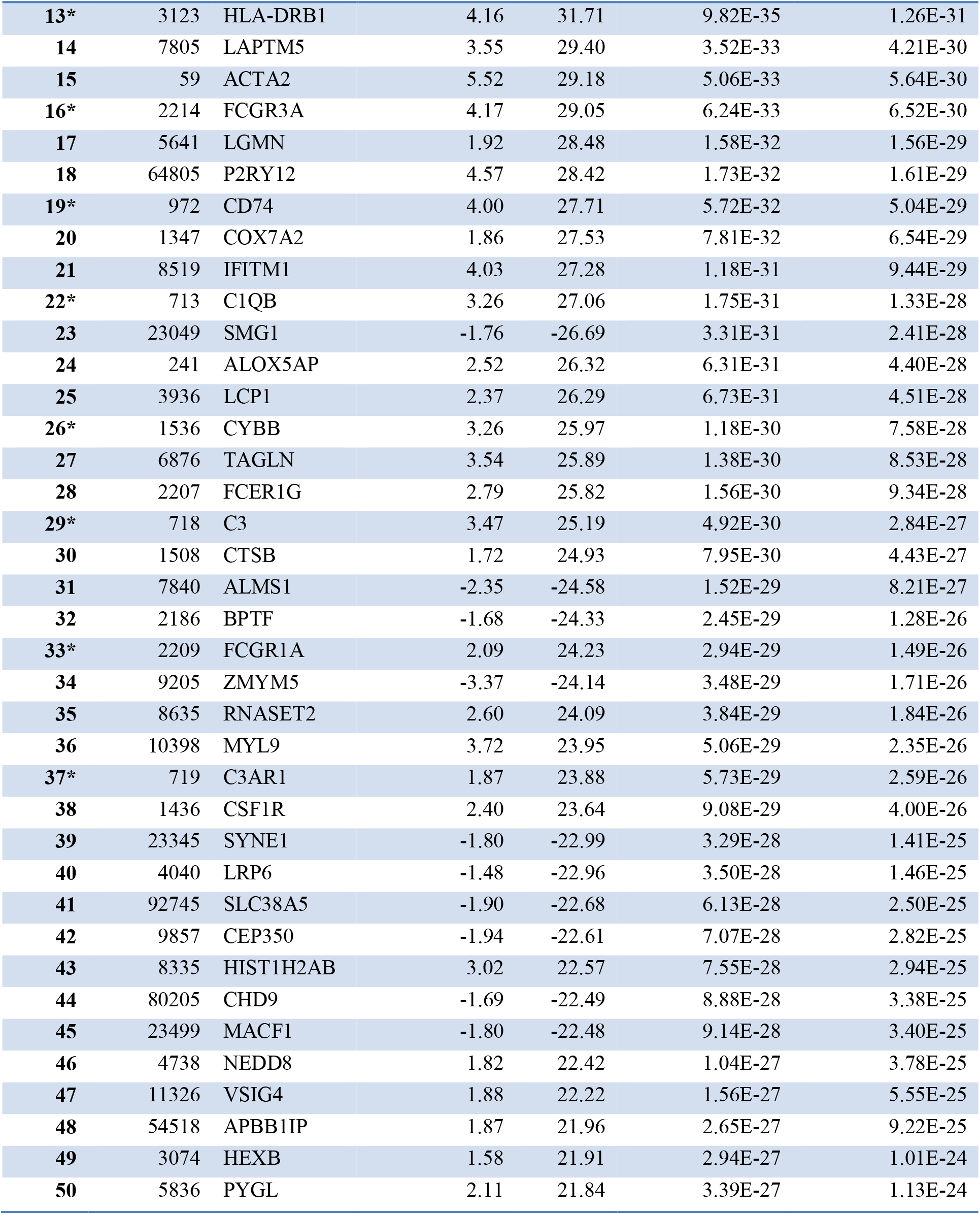
Top 50 differentially expressed genes in male versus female at prenatal Stage 7 (before birth) cortical development. Ranking is based on the adjusted p–value (top=lowest p–value). Genes that are discussed in the text are marked with an asterisk.

1. *Complement components*: C1QB (Ranked #22, logFC=3.26, p-value= 1.33E-28), C3 (Ranked #29, logFC= 3.47, p-value= 2.84E-27), C3AR1 (Ranked # 37, logFC= 1.87, p-value= 2.6E-26)
2. *Fc receptors*: FCGR3A (Ranked #16, logFC= 4.17, p-value= 6.52E-30), FCGR1A (Ranked #33, logFC= 2.1, p-value= 1.5E-26),
3. *Members of the nicotinamide adenine dinucleotide phosphate (NADPH) oxidase complex*, gp91: CYBB (Ranked #26, logFC=3.26, p-value= 7.58E-28),
4. *Toll-like receptors*: CD14 (Ranked # 12, logFC=4.52, p-value= 3E-32),
5. *Major histocompatibility complex molecules:* HLA-DRB1 (Ranked # 13, logFC=4.16, p-value= 1.26E-31), CD74 (Ranked # 19, logFC=4, p-value=5.04E-29).

An enrichment analysis restricted to KEGG pathways showed that inflammatory enriched pathways that were enriched within the 50 top differentially expressed genes between male and female at Stage 7 fell principally into five groups (complement cascade, Fc receptors, NADPH-oxidase complex, major histocompatibility complex molecules and Toll-like receptors). The convergence across these five groups is best captured by “Complement and Coagulation Cascade” and “Phagosome” KEGG pathways as it is shown in Supplementary File 3.

Figure 3 shows microglia-mediated synaptic pruning and tagging genes [20, 22, 29, 30] highlighting a common pattern of sex differential expression for CXCR1, C1QC, C1QB, C3 at Stages 7 (before birth, more highly expressed in males) and 9 (after birth, more highly expressed in females). Another male-biased stage was Stage 11 (middle and late childhood) with complement genes C1QC, C1QB and C3. CX3C chemokine receptor 1 (CX3CR1) on microglia and its ligand, CX3CL1, signaling is required for the migration of sufficient numbers of microglia into the brain and for microglia-mediated synaptic pruning in early development and spine formation and elimination in mature circuits. C1Q initiates the complement cascade, leading to cleavage of C3, which binds to synaptic surfaces for microglial recognition and subsequent pruning [30].

**Figure 3:**
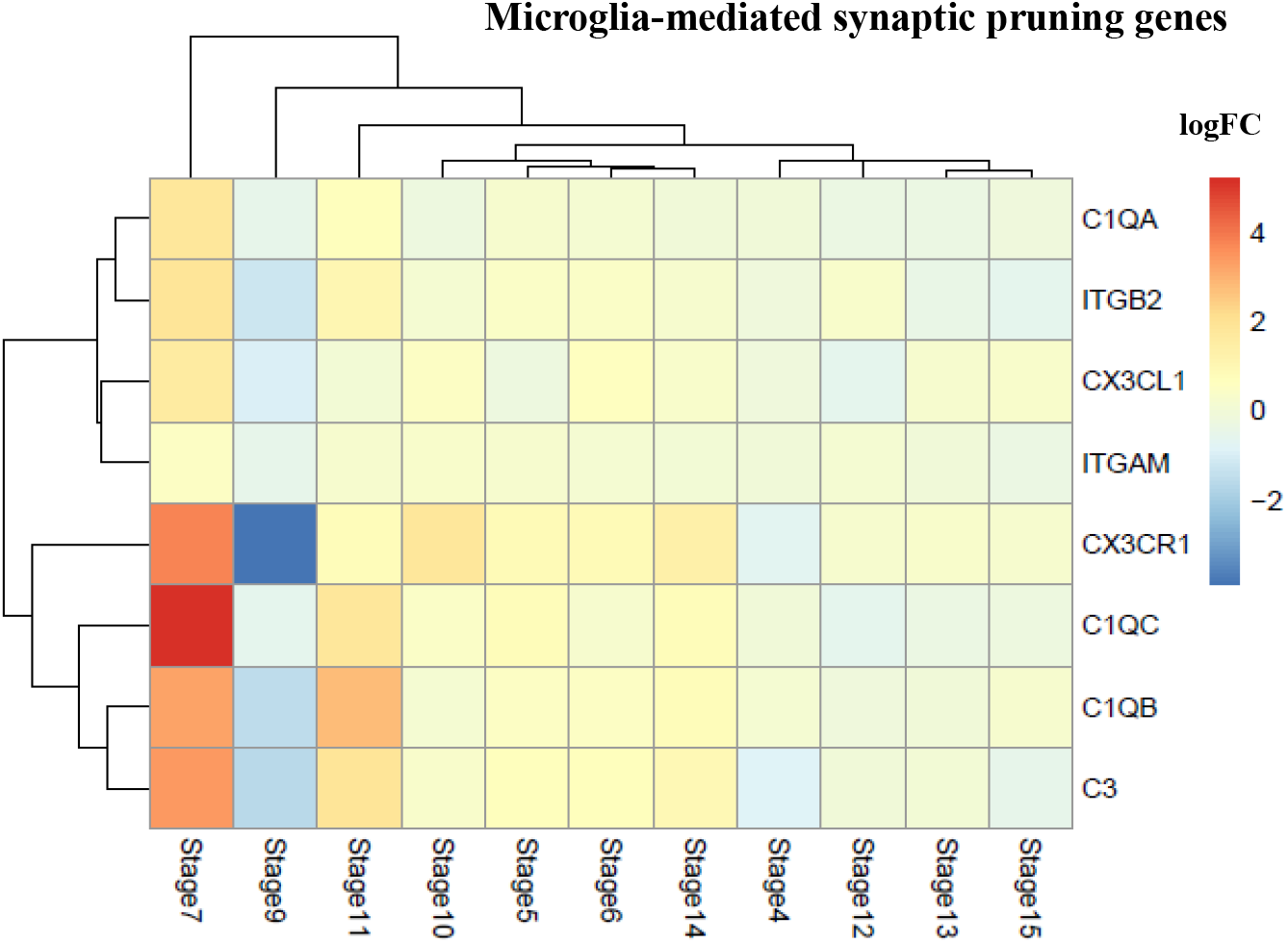
Hierarchical clustering of logFC (Fold Change=males/females) of expression of microglia-mediated synaptic pruning genes in males versus females. Red=Upregulated in males versus females; Blue=Upregulated in females versus males; Stage 7=Before birth; Stage 9=After birth.

The small number of samples at each stage of development is a limiting factor in the current analysis. However, the signal appears robust under various analyses including combining Stage 6 and 7 (Supplementary File 4).

## Discussion and Conclusions

We hypothesized that immunological signaling play a key role in underwriting the sexual dimorphism of the developing cortex and the subsequent development of some neurological disorders that have a strong sex bias in their symptoms or prevalence. We focused on microglial processes and the complement cascade-dependent synaptic pruning/tagging. Specifically, we sought and found evidence of a sexually dimorphic developmentally regulated module that includes immune activation of microglia.

The question then arises whether this developmental sexually dimorphic model is related to human disease. Males are more likely to be diagnosed with neurological disorders that have developmental origins such as autism that tend to occur early in life. Females are more likely to be diagnosed with neurological disorders that tend to appear later in life or after puberty such as anxiety disorders and depression. Do these differences in disease prevalence at different ages relate to increased prenatal vulnerability in males to immune activation and, conversely, increased vulnerability of females to postnatal immune activation? Bilbo et al., investigated various aspects of these questions [1] and recently identified a microglia-specific gene expression program in mice that was used to create a microglia developmental index. This index was applied to reveal differences between males and females. They showed that the gene expression program was <u>delayed</u> in males relative to females and exposure of adult male mice to LPS, a potent immune activator, accelerated microglial development in males [27]. They also found that brain regions involved in cognition, learning and memory, such as hippocampus, parietal cortex and the amygdala have significantly more microglia in males than females in the postnatal rodent brain [1, 26]. Microglia produce high levels of cytokines and chemokines [26], which might lead to a higher concentration of inflammatory processes contributing to an “inflammatory load” on male brain, perturbation of normal organ/tissue function and therefore easier susceptibility to develop “dysregulated phenotype”. In addition, male rats infected with *E.coli* on postnatal day 4 have long-term changes in the function of microglia [31–33], while the same immune challenge at postnatal day 30 has no long-term effects on glial function or behavior into adulthood [34]. On the other hand, no long term effects on cognitive behavior were observed in female rats treated similarly [35].

In our analysis, we identified significant gene expression and KEGG pathway differences in developing human male versus female cortex that were related to immune processes and infections (i.e., immune activation) and to neurological diseases, especially at the stages before birth, puberty and late adulthood (Figure 1). Specifically, we have shown that the male brain is enriched for the expression of genes associated with phagocytic function of microglia through complement-dependent synaptic pruning especially before birth (Figures 2 and 3). One mechanism involved in both neurodevelopment and models of neurodegeneration is the classical complement cascade, which is activated when microglia target synapses or other unwanted neural material for phagocytosis [16, 17, 20, 22, 36]. Stevens et al. have shown that C1q, the initiating protein of the cascade, and C3, a downstream protein, localize to subsets of immature synapses, likely marking them for elimination. Microglia, which express CR3, engulf these synaptic terminals through the C3-CR3 signaling pathway [17, 29]. Besides the complement cascade, another signaling pathway that is critical for synaptic pruning is the microglial fractalkine receptor (CX3CR1). Our analysis shows that all the major components in synaptic pruning by microglia are upregulated in males versus females in the developing cortex at the stage before birth (prenatal stage), and are downregulated in males versus females at early infancy (postnatal stage).

In addition to the classical complement cascade, we found Fc receptors upregulated in males. Fc receptors are generally used in the periphery for a classical process of pathogen opsonization (process of identifying invading particle to phagocyte) and subsequent phagocytosis. The systems involved in peripheral opsonization include the complement cascade (C3b/C4b) and Fc receptor/IgG. Unlike the complement cascade, Fc/IgG has not been fully investigated for its role in synaptic tagging/pruning because the physical size of immunoglobulins limits their ability to cross the brain-blood barrier. Nonetheless, there is growing evidence for the increased expression of Fc gamma receptors in resident CNS cells including microglia and neurons during aging, and their involvement in the pathogenesis of age-related neurodegenerative diseases [37]. Systemic inflammation has also been shown to lead to increased serum-derived IgG expression in the brain parenchyma, suggesting a role for IgG - Fc receptor interaction in switching primed microglia to an aggressive pro-inflammatory phenotype [38]. Our results suggest that Fc receptors might be also linked to microglial synaptic phagocytosis in brain or immune signaling in brain in general.

In summary, our results suggest that microglial function is sexually dimorphic at different stages of neurodevelopment. We identified a common set of molecular pathways that, in males, is involved in prenatal immune activation which might render males more susceptible to developing some neurological disorders, while in females, is involved in postnatal immune activation which might protect females from the same disorders. This could explain the possible sexual dimorphic comorbidities of specific autoimmune disorders and neurodegenerative conditions. Our results are limited by the sample size of our primary dataset. More definitive evidence would involve measuring the corresponding protein expressions of the candidate pathways (complement cascade, Fc receptors) in a greater number of independent human brain tissue samples.

